# AmyloCore-ML: AI/Machine Learning Enabled Identification of Amyloid Fibril Core Regions Using Protein Language Models

**DOI:** 10.64898/2026.09.25.754455

**Authors:** Jasdeep Singh, Raunak Jangid, Ankit Srivastava

## Abstract

Identifying protein regions incorporated into amyloid fibril cores is important for understanding protein aggregation and its involvement in associated neurodegenerative and systemic disorders. Existing sequence-based predictors have largely focused on identifying short aggregation-prone motifs, which do not necessarily correspond to the extended regions forming experimentally resolved amyloid fibril cores. Here, we introduce AmyloCore-ML, a machine-learning framework designed to predict fibril-core regions directly from protein sequences. Using experimentally determined amyloid fibril structures, we curated a dataset comprising 69 proteins and 82 independent amyloid-core regions, with structurally assigned core residues and matched non-core residues derived from the same parent proteins. Sequence regions were encoded using physicochemical descriptors and protein language model (PLM) embeddings, including ESM-2, ANKH, and ProtT5, and evaluated through protein-grouped cross-validation. PLM-derived embeddings consistently outperformed physicochemical descriptors. The best-performing model, combining ESM-2 embeddings with an ExtraTrees classifier, achieved an AUROC of 0.833 and a mean peak distance of 12.4 residues, while demonstrating improved performance against selected existing amyloid predictors. AmyloCore-ML is easily accessible through an interactive Google Colab (https://github.com/jasdeep002/AmyloCore-ML), enabling sequence-based amyloid fibril-core prediction without local installation or dedicated computing infrastructure.

## Introduction

Protein aggregation into amyloid fibrils is associated with a broad spectrum of neurodegenerative and systemic disorders, including Alzheimer’s disease (AD), Parkinson’s disease, tauopathies, transthyretin amyloidosis and prion diseases^1-3^. Irrespective of protein sequence, native structure or biological function, mature amyloid fibrils forms a beta-rich architecture in which beta-strands run approximately perpendicular to the fibril axis^1, 4^. The mature fibrils in various diseases, however do not share structural similarity^5, 6^. Addtionally, full protein sequence may not participate in amyloid formation. In many experimentally determined fibril structures, only certain regions become incorporated into the highly ordered fibril core, whereas the remaining sequence remains conformationally heterogeneous as non-core regions^4^. Predicting the structurally ordered regions directly from sequence therefore represents an important gap in amyloid biology.

A broad range of computational methods has been developed to identify aggregation-prone regions in proteins, which could be an entire core region or fraction of it. Current sequence-based predictors, including TANGO^7^, WALTZ^8^, Zyggregator^9^ and AGGRESCAN^10, 11^, use physicochemical properties, beta-sheet propensity, residue aggregation scales or sequence-window statistics to identify local regions with elevated aggregation potential. Structure-informed methods additionally account for conformational context and solvent accessibility^12^. Most recently, AggrescanAI demonstrated that contextual embeddings from PLMs can support residue-level aggregation prediction directly from primary sequence^13^. These approaches have substantially advanced the prediction of aggregation propensity. However, an aggregation-prone region and the structurally ordered core of an amyloid fibril represent related but distinct properties. Short peptides may display strong intrinsic amyloidogenicity when isolated experimentally but may or may not form the ordered core of the corresponding full-length protein^14-17^. For example Tau protein contains two aggregation-prone regions: ^275^VQIINK^280^ and ^306^VQIVYK^311^; however, in its AD fibrils, the structurally ordered amyloid core is substantially larger, extending approximately from V306 to F378. Similarly, residues outside an experimentally resolved fibril core may influence nucleation or fibril formation without remaining ordered in the mature structure^18-20^. Consequently, predictors trained primarily using experimentally annotated aggregation-prone peptides or broader amyloidogenic regions are not explicitly optimized to identify the sequence that is ultimately incorporated into a resolved amyloid structure.

The identification of disease associated fibrils using cryo-EM or solid-state structural studies provides an opportunity to address this distinction directly^21^. Using high resolution fibril structures from Amyloid Atlas 2026^21^, we obtained residue-level structural annotations to obtain amyloid-core localization. For each amyloid-forming protein, sequences outside the experimentally ordered core were used as matched non-core controls. This differs from previous classification strategies in which amyloidogenic sequences and negatives are obtained from separate protein populations and therefore reduces protein-level differences unrelated to local core formation. Thus, the predictor learns the sequence grammar of regions that become structurally incorporated into a fibril under a given biochemical or disease condition, from non-core regions of the same amyloid-capable protein.

Leveraging the sequence grammar of known amyloid forming proteins, we present a sequence-based predictor for amyloid fibril core localization-AmyloCore-ML. Experimentally defined core and matched non-core regions were converted into multiple sequence-window representations and encoded using conventional physicochemical descriptors and contextual protein language model embeddings, namely ESM-2 (Evolutionary Scale Modeling-2)^22^, Ankh^23^ and ProtT5^24^. AmyloCore-ML produces a continuous positional core-propensity profile that can be mapped directly onto an input protein sequence. This study provides a structure-guided, sequence-based platform for predicting amyloid core-forming regions and defining the sequence features associated with their incorporation into fibrils. Such predictions may be particularly useful for the rational design of proteomimetics or other inhibitors intended to engage extended fibril-core surfaces rather than isolated short aggregation-prone motifs. An overview of the prediction workflow is presented in Figure 1.

**Figure 1.**
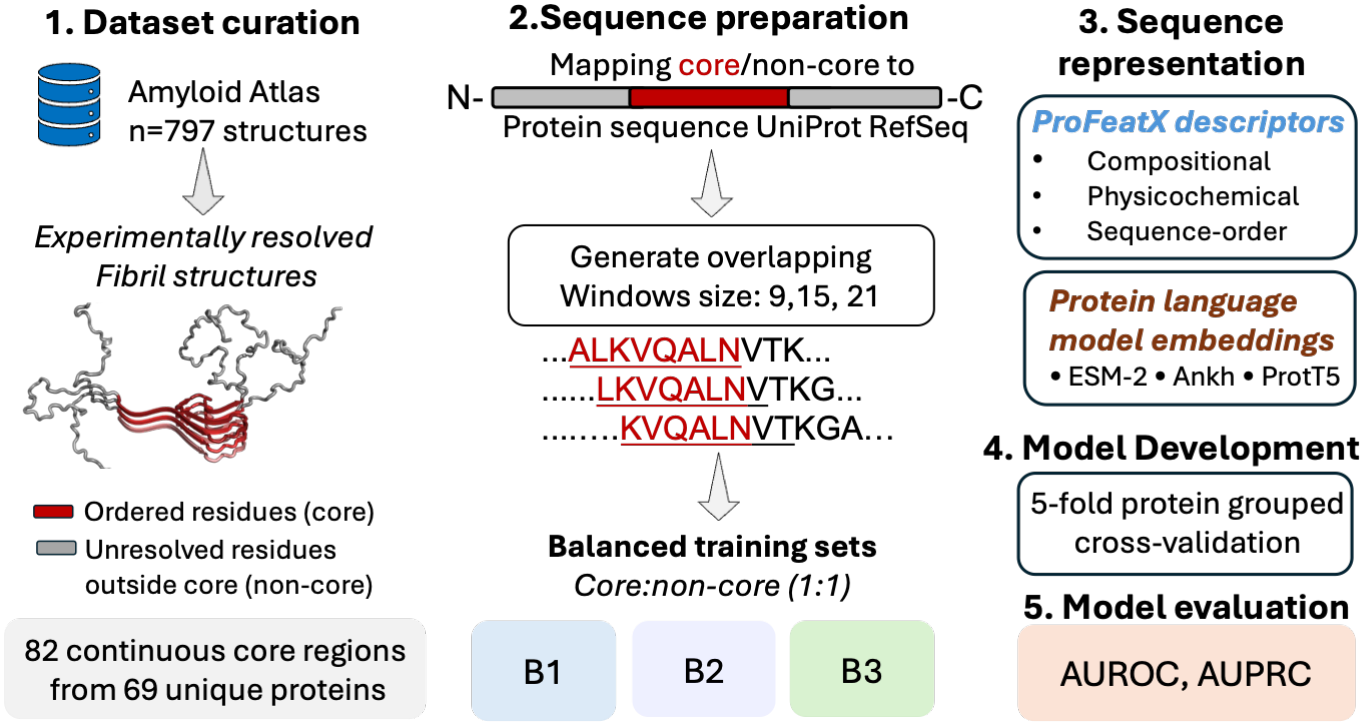
Structure-informed framework for amyloid fibril-core prediction. Experimentally resolved amyloid structures were mapped to full-length protein sequences, yielding 82 continuous core regions from 69 proteins. Core and non-core regions were evaluated across three benchmarking schemes (B1-B3) using ProFeatX descriptors, protein language-model embeddings, and multimodal models. These representations were subsequently used for machine-learning analysis, including individual and integrated/multimodal models. Predictive performance was quantified primarily using the area under the receiver-operating characteristic curve (AUROC) and area under the precision-recall curve (AUPRC).

## Methods

### Structural dataset and definition of core regions

Experimentally resolved amyloid fibril structures were obtained through Amyloid Atlas (n=797 entries as of April 2026)^21^ (Figure 2a). De novo designed amyloids and synthetic non-biological assemblies were excluded, and structural annotations were checked for redundancy and against corresponding reference sequences obtained from UniProt^25^. The non-redundant dataset comprised 82 continuous amyloid regions corresponding to 69 unique proteins (Figure 2b and Supplementary Sheet 1). Pairwise sequence similarity among the 69 proteins was assessed using all-versus-all global protein sequence alignment using MMseqs2 with the BLOSUM62 substitution matrix and affine gap penalties^26^. Full-length sequence identity was calculated as the proportion of identical residues across the complete global alignment. For discontinuous ordered regions within the same protein, each core region was retained. The resulting dataset was divided into two sequence classes. Residues experimentally observed within the ordered fibril structure were designated core, whereas eligible residues outside all annotated structural cores were designated non-core. At the residue level, the final dataset contained 4,819 unique core residues and 10,433 non-core residues (Figure 2c).

**Figure 2.**
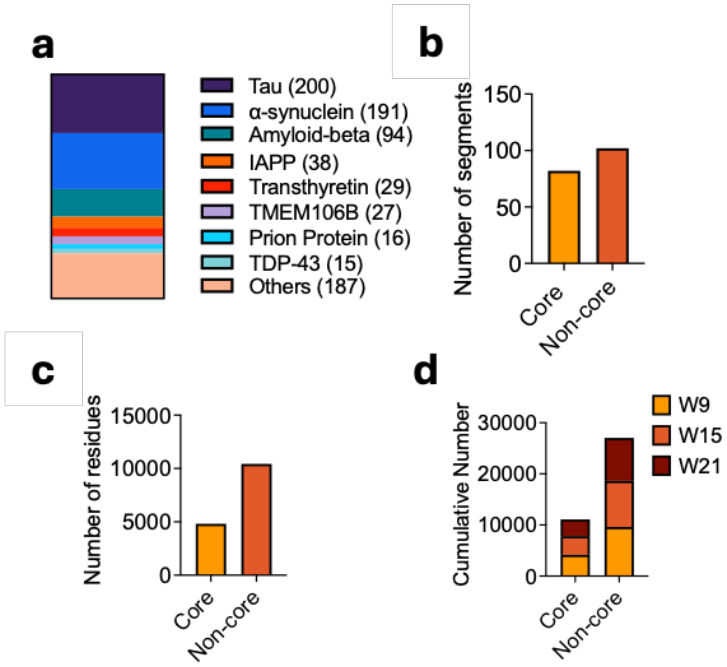
Dataset composition. **(a)** Distribution of amyloid proteins represented in the curated structural dataset obtained from Amyloid Atlas **(b-c)** Number of core/non-core segments and corresponding residues. **(d)** Cumulative number of sequence windows generated using 9-, 15-, and 21-residue windows (W9, W15, W21).

### Generation of sequence windows

Core and non-core regions were divided into overlapping windows of 9, 15, and 21 residues (step size of 1 and without padding), thus preserving class boundaries. The resulting datasets: W9 contained 13,780 windows (4,163 core; 9,617 non-core) from 69 proteins, W15 contained 12,716 (3,684 core; 9,032 non-core) from 68 proteins, and W21 contained 11,730 (3,240 core; 8,490 non-core) from 65 proteins (Figure 2d). To address class imbalance and bias from randomly selected non-core windows, three balanced datasets (B1, B2, and B3) were generated independently for each window size with 1:1 core:non-core ratios. Non-core overlap among balanced replicates ranged from 34% for W9 to 18-20% for W21.

### Sequence representations

Two complementary classes of sequence representation were evaluated. First, classical descriptors were calculated to represent amino-acid composition, physicochemical characteristics, and sequence-order information using ProFeatX^27^. Features requiring external predictions of secondary structure, solvent accessibility, disorder, or related structural properties were excluded. Second, contextual embeddings were also generated using ESM-2, Ankh, and ProtT5^22-24^. The same window sequences and protein partitions were used for every representation, allowing direct comparison under identical conditions. Multiple classifiers were selected and screened for comparative model performance. Hyperparameter optimization and, where applicable, feature selection were restricted to training proteins.

### Protein-grouped cross-validation was implemented to prevent sequence leakage

To prevent sequence leakage from W9, W15, and W21 sliding windows, a single master five-fold protein-grouped cross-validation scheme was generated across the 69 proteins. All windows from a given protein were assigned to the same outer fold, and these were also retained across B1, B2, and B3 datasets. Feature ranking was performed separately within each outer fold using only training proteins. Model performance was evaluated independently for each held-out protein and then summarized across proteins. AUROC and AUPRC were used to measure residue-ranking performance, while localization was assessed using distance of the highest-scoring region from the experimental core.

### Full-length prediction and residue reconstruction

For full-length prediction, every possible window of the selected size was generated using a stride of one residue. Each window was embedded using the corresponding protein language model and assigned a core probability. A residue generally occurs in several overlapping windows. Its final probability, P_core, was therefore defined as the mean probability of all windows containing that residue. This reconstruction produces a continuous probability profile along the complete protein sequence. A threshold of P**_**core ≥ 0.50 was retained for segment-level analyses. Threshold-independent AUROC and AUPRC were evaluated separately so that ranking performance was not conflated with probability calibration. For benchmarking, AmyloCore-ML predictions were compared directly against AggrescanAI and CrossBeta predictors. AggrescanAI uses deep learning with contextual ProtT5 protein-language-model embeddings to generate residue-level aggregation-propensity predictions, whereas the Cross-Beta predictor uses an ExtraTrees machine-learning framework trained on experimentally characterized cross-β amyloid regions. Together, these methods provide comparison with both protein-language-model-based and conventional machine-learning approaches developed for amyloidogenic-region prediction. For the unseen evaluation, we used four proteins that were not present in AmyloidAtlas and were excluded from model development: horse myoglobin(Uniprot:P68082, PDB: 27WR), P02070 (PDB: 20YJ), Oat 11S globulin (Uniprot: Q38779, PDB: 26PC), Oat 12S seed storage globulin 2 (UniProt: P14812, PDB: 9VAM), and human Integral membrane protein 2B (UniProt: Q9Y287, PDB: 9OI9). Predictions were generated using the locked AmyloCore-ML models without retraining or parameter adjustment, and predicted CORE regions were compared directly with the experimentally resolved fibril-core boundaries from the corresponding structures.

### Final predictor

Cross-validation was used for model comparison and performance estimation. After the representation and model design were fixed, production models were refitted using all available development proteins. Fresh ESM-2 and Ankh embeddings were generated for the originally labelled W9, W15, and W21 windows. Separate classifiers were trained for B1, B2, and B3. During prediction, probabilities from the three classifiers were averaged before reconstruction of the residue-level profile. The final implementation as AmyloCore-ML allows the user to select ESM-2 or Ankh and W9, W15, or W21, or visualize all six profiles simultaneously. The output consists of residue-level P_core values, predicted core segments, and graphical profiles along the protein sequences.

## Results

We carried out this study to determine whether using sequence-features only, if we could identify and distinguish structurally resolved amyloid fibril-core regions from matched non-core regions within proteins. Across 2,346 unique protein pairs generated from 69 proteins, global sequence identity was generally low, with a mean of 12.72% and median of 11.33%. 17 pairs (0.51%) showed ≥70% identity, indicating limited full-length sequence redundancy across the dataset (Figure S1). To build an optimal predictor, we systematically compared physicochemical (ProFeatX) descriptors and protein-language-model embeddings-ESM-2, ProtT5, and ANKH across three sequence window sizes (W9, W15, and W21), three independently balanced core/non-core datasets (B1-B3), using a range of classifiers. All configurations were evaluated using the same protein-grouped cross-validation framework, allowing the effects of sequence representation, window size, balancing strategy, and classifier choice to be compared directly while preventing windows from the same protein from being split across training and validation sets. Firstly, we noticed that performance was comparatively invariable across non-core datasets (B1, B2 and B3), with overall mean AUROC and AUPRC ranging from 0.763 to 0.814 and 0.7742 to 0.801 respectively, arguing against dependence on a particular negative sample (Figure 3 and Figure S2). However, we observed that Sequence context (window size) did have an effect on performance. Averaged across classifiers, AUROC increased from 0.755 at W9 to 0.820 at W21 for ProFeatX, whereas the corresponding increases were 0.769-0.867 for Ankh, 0.768-0.851 for ESM-2, and 0.774-0.854 for ProtT5 (Figure S2). Specifically at W21, ExtraTrees achieved AUROC/AUPRC values of 0.875/0.848 with Ankh, 0.869/0.841 with ESM-2, and 0.868/0.842 with ProtT5. RBF-SVC (Radial Basis Function-Support Vector Classifier) also produced the same trend, reaching 0.871/0.843 with Ankh (Figure 3). The agreement between tree-based and nonlinear kernel classifiers indicates that the classification metrics were not specific to one decision architecture, while the improvement over physicochemical descriptors suggests that contextual sequence relationships captured by PLMs (ESM-2, Ankh, ProtT5) could provide additional information relevant to structural core formation.

**Figure 3.**
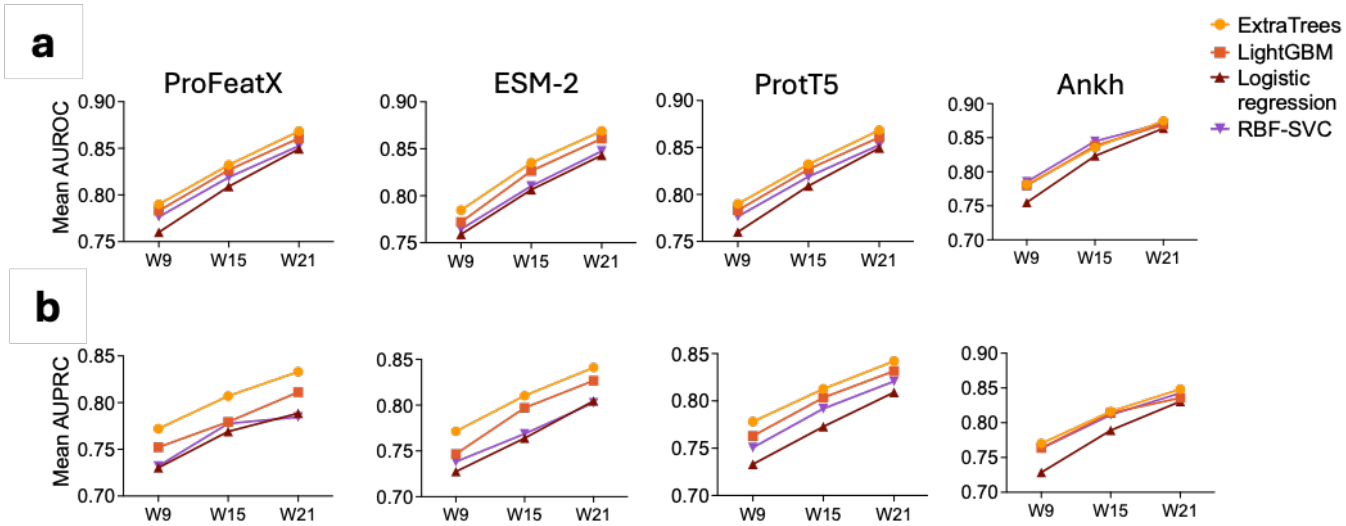
Cross-validated discrimination of experimentally resolved amyloid CORE and matched NONCORE windows. **(a-b)** Mean AUROC and AUPRC, respectively, for ExtraTrees, LightGBM, Logistic regression, and RBF-SVC models using ProFeatX, ESM-2, ProtT5, and Ankh representations across the three window sizes.

Next, we asked whether ExtraTrees and RBF-SVC, top-performers obtained from training on separate core and non-core regions, also carry localization potential in complete protein sequences, without pre-assigning core or non-core information. Across representations and both classifiers, mean residue-level AUROC increased from 0.789 at W9 to 0.860 at W21, accompanied by an increase in AUPRC from 0.686 to 0.758. The mean distance between the predicted probability peak and the experimentally defined amyloid-core region also decreased with increasing window length for ExtraTrees, from approximately 15 residues at W9 to 11 residues at W21(Figure S3). ExtraTrees and RBF-SVC displayed complementary behavior: RBF-SVC produced greater core recovery and segment overlap, whereas ExtraTrees provided stronger residue ranking and precision. This complementarity motivated retention of both classifiers in the locked-model analysis rather than selection of a single algorithm.

Finally, locked models were subsequently applied to full-length proteins without retraining and compared against two established sequence-based amyloid predictors, CrossBeta and AggrescanAI, using the same experimentally defined core annotations (Figure 4). ESM-2/ExtraTrees W21 provided a particularly strong balance between discrimination and localization, with a residue AUROC of 0.833, AUPRC of 0.751, and mean peak distance of 12.4 residues. Ankh/RBF-SVC W15 produced similarly strong performance (AUROC 0.824; AUPRC 0.739) with a mean peak distance of 14.4 residues and recovered 70.4% of core residues at *P*_*CORE*_ ≥ 0.50. Notably, the highest AUROC alone did not identify the best-localizing model: ProtT5/ExtraTrees W21 reached an AUROC of 0.838 but had a mean peak distance of 24.1 residues. CrossBeta achieved a mean residue-level AUROC of 0.718 and AUPRC of 0.659, with a mean peak distance of 22.7 residues. AggrescanAI produced a similar AUROC of 0.719 but a lower AUPRC of 0.621 and a larger mean peak distance of 30.2 residues (Figure 4c). Collectively, the ESM-2/ExtraTrees W21 and Ankh/RBF-SVC W15 models combined higher residue-ranking performance with shorter mean distances to the experimentally resolved core. Lastly, in the independent unseen benchmark, both ESM-2 and ANKH models identified high-probability core segments that overlapped or localized near the experimentally resolved fibril-core regions across several proteins (Figure 4d). The consistency of these predictions across multiple window sizes supports the ability of the AmyloCore-ML to generalize core localization beyond the proteins used for model development. An interactive interface for the AmyloCore-ML Predictor is provided as Figure S4. Users can provide protein sequence(s) and subsequently select sequence representation, window size, core probability threshold to generate residue level amyloid core predictions.

**Figure 4.**
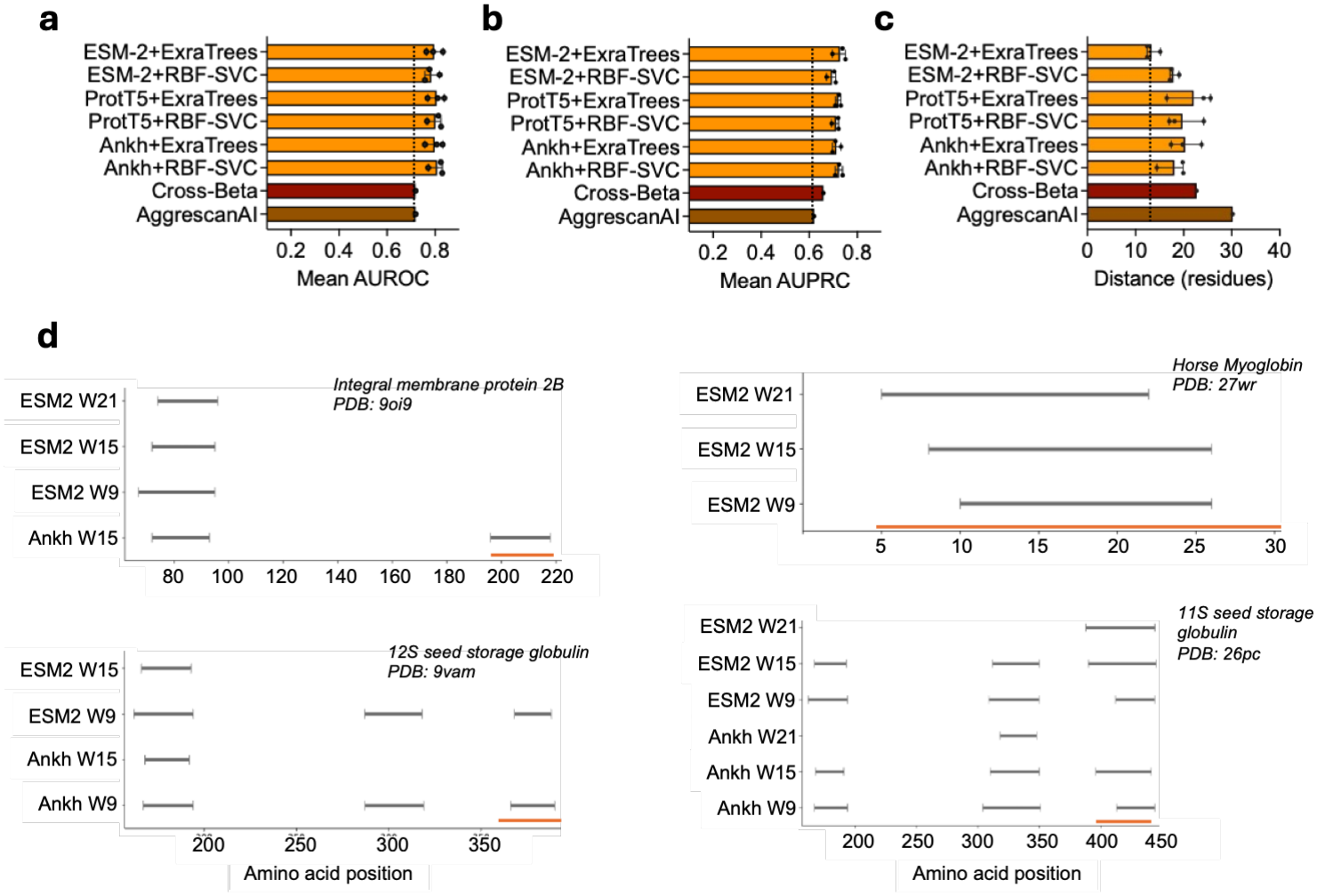
Benchmarking of amyloid-core prediction methods. Comparison of ESM-2, ProtT5, and Ankh embeddings with ExtraTrees and RBF-SVC classifiers against Cross-Beta and AggrescanAI. **(a)** Mean AUROC, **(b)** mean AUPRC, and **(c)** mean distance between the predicted and experimentally defined amyloid-core regions. Error bars from symbols indicate variability across three windows: W9, W15, W21. (d) Predicted core segments from locked ESM-2 and ANKH models are shown for different sequence-window sizes (W9, W15, and W21) across representative proteins from the unseen benchmark dataset. Gray horizontal bars indicate predicted core regions, whereas orange bars denote the experimentally resolved fibril-core region derived from the corresponding PDB structure.

## Discussion

In current work, a sequence-based framework for predicting regions preferentially incorporated into experimentally resolved amyloid fibril cores is presented. The approach builds on complementary developments in amyloid prediction, including experimentally validated aggregation-prone regions, structure-based compatibility models, machine-learning classifiers, and contextual protein language representations^9, 11-13^. A central feature of the present framework is the use of structure-defined fibril cores together with matched non-core sequences from the same amyloid-forming proteins. This shifts the prediction problem from general amyloidogenicity towards-identifying which regions of an amyloid-forming protein are preferentially incorporated into the ordered fibril architecture.

The curated dataset comprises 69 proteins, 82 independent amyloid core regions. In our work, longer sequence windows generally showed higher AUROC/AUPRC, with the strongest cross-validated models. This trend suggests that structural-core identity is not encoded exclusively by short amyloidogenic motifs, but also depends on the surrounding sequence environment. From the predictor standpoint, our design complements previous approaches using experimentally characterized peptides, literature-annotated APRs, or structurally defined amyloid sequences. For example, Cross-Beta was trained on naturally occurring cross-β amyloid regions using engineered features, with ExtraTrees selected as the final classifier^28^. AggrescanAI, for instance, represents a closely related methodological advance as it uses ProtT5 contextual embeddings and deep learning to predict residue-level aggregation-prone regions directly from sequence ^13^; however, its training endpoint is experimentally annotated APRs rather than residues specifically resolved within mature fibril structures. In our structural-core benchmark, both methods showed lower residue-level discrimination and larger mean peak distances than the strongest locked PLM models (Figure 4). However, these comparisons should be interpreted in the context of different training objectives: AggrescanAI and CrossBeta were developed to identify aggregation- or cross-β-prone sequence regions, whereas the present models were explicitly trained against residues incorporated into experimentally resolved mature fibril cores. This distinction defines the biological endpoint of the framework. Because core and non-core sequences were sampled from the same amyloid-forming proteins, broad protein-level properties alone cannot readily explain their separation. Instead, the models are directed toward local sequence features associated with incorporation into the ordered fibril structure. The systematic benefit of longer windows further indicates that this information extends beyond isolated short amyloidogenic motifs and is distributed over a broader local sequence environment. Accordingly, *P*_*CORE*_ should be interpreted as structural amyloid-core propensity rather than absolute aggregation propensity. Non-core residues may still contribute to nucleation, fibril stability, or alternative polymorphs without remaining ordered in a particular mature fibril^29, 30^. Nevertheless, concordance across independent non-core samplings, physicochemical and language-model representations, classifiers, and external predictor benchmarking provides convergent evidence that experimentally resolved amyloid cores contain a learnable and spatially recoverable sequence signature.

## Supporting information

indicating limited full-length sequence redundancy across the dataset

The non-redundant dataset comprised 82 continuous amyloid regions corresponding to 69 unique proteins

## CRediT authorship contribution statement

JS: Conceptualization, Methodology, Investigation, Formal analysis, Resources, Supervision, Project administration, Data curation, Visualization, Writing - original draft, Writing - review & editing, Project administration, Supervision. RJ: Methodology, Investigation, Formal analysis, Data curation, Validation, Writing - review & editing. AK: Methodology, Formal analysis, Resources, Writing - review & editing.

