## Supplementary material for "AmyloCore-ML: AI/Machine Learning Enabled Identification of Amyloid Fibril Core Regions Using Protein Language Models": indicating limited full-length sequence redundancy across the dataset

Jasdeep Singh[1#], Raunak Jangid[2], Ankit Srivastava[2,3#]
[1] Department of Chemistry and Biochemistry, University of Denver, Colorado, USA
[2] Department of Biotechnology and Chemical Engineering, Manipal University Jaipur, India
[3] Center for Health and Therapeutics Research (CHTR), Manipal University Jaipur, India

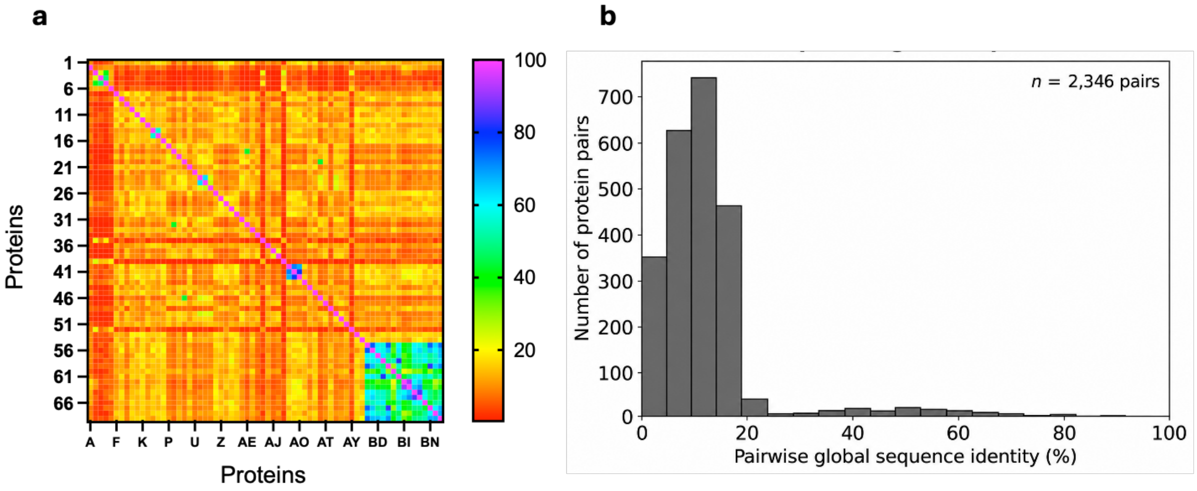


**Figure S1. Global sequence-identity analysis of the curated amyloid protein dataset. (A)** Pairwise global sequence-identity matrix for the 69 proteins, with the diagonal representing self-identity and off-diagonal values indicating full-length pairwise sequence identity. **(B)** Distribution of the 2,346 unique non-self pairwise global sequence identities, showing that most protein pairs exhibit low sequence identity, with only a small fraction extending into higher-identity ranges.

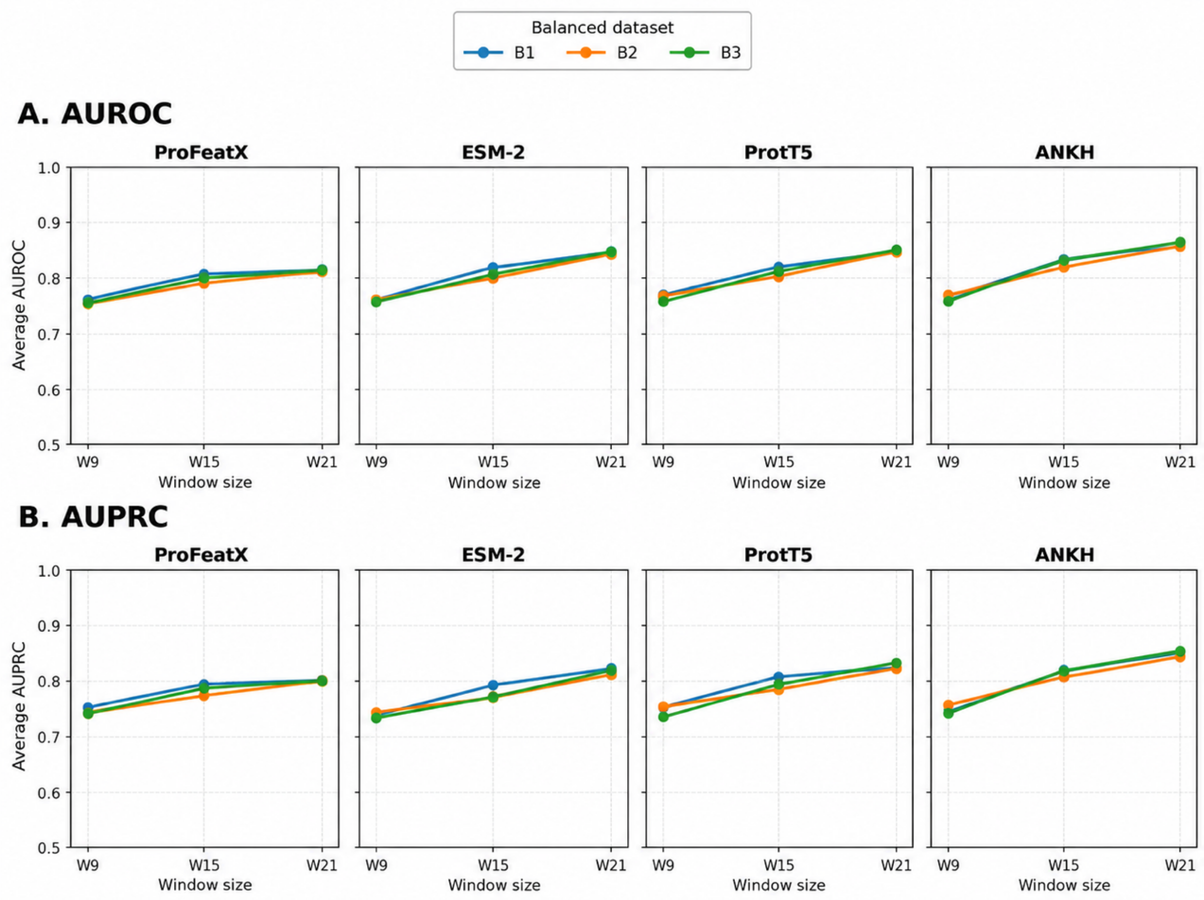


Figure S2. **Performance metrics across sequence window size and dataset balancing strategy on protein representations.** Pooled out-of-fold performance was compared for ProFeatX descriptors and protein language model embeddings from ESM-2, ProtT5, and ANKH using window sizes of 9, 15, and 21 residues. (A) Mean AUROC and (B) mean AUPRC across the three balanced datasets (B1–B3). Values represent averages across the four classifiers used consistently across all representations: ExtraTrees, LightGBM, logistic regression, and SVC.

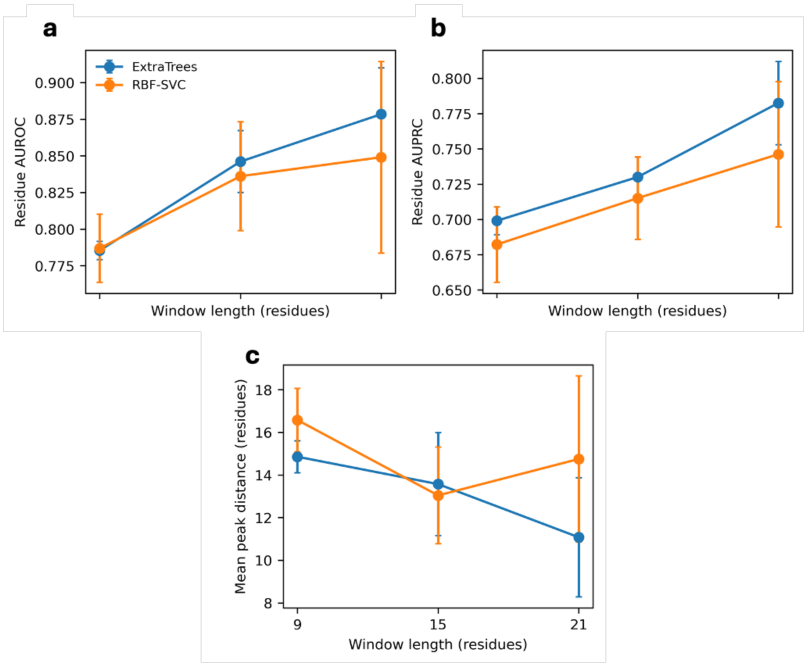


**Figure S3. Sequence context and classifier architecture influence full-length CORE localization.** Performance of the ExtraTrees (Extremely Randomized Trees) and RBF-SVC models was evaluated across sequence windows of 9, 15, and 21 residues on full-length proteins (a) Residue-level AUROC and (b) residue-level AUPRC across window lengths. (c) Mean distance, in residues, between the predicted probability peak and the experimentally defined amyloid-core region. Points represent mean performance and error bars indicate variability across evaluated proteins.

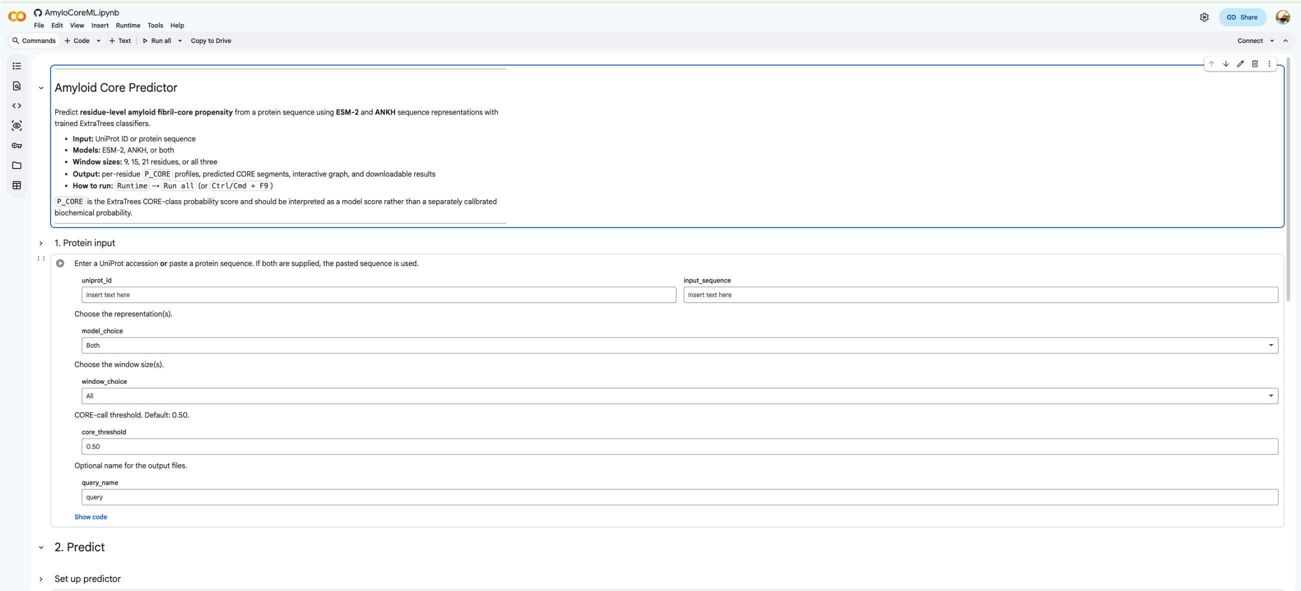


**Figure S4**. **Interactive Google Colab interface for the AmyloCore-ML Predictor.** The notebook provides a user-accessible implementation of the prediction framework for identifying residue-level amyloid fibril-core propensity from protein sequence. Users can supply a UniProt accession or amino acid sequence, select the sequence representation and window size, define the CORE-call threshold, and assign an output query name. The interface supports prediction using ESM-2 and ANKH-based models and generates residue-level prediction outputs for downstream visualization and analysis.
